# Focus Your Screening Library: Rapid Identification of Novel PDE2 Inhibitors with *in silico* Driven Library Prioritization and MicroScale Thermophoresis

**DOI:** 10.1101/2020.04.22.021360

**Authors:** Florian Kaiser, Maximilian G. Plach, Thomas Schubert, V. Joachim Haupt

**Affiliations:** 2bind GmbH, Regensburg, Germany | 2bind.com; PharmAI GmbH, Dresden, Germany | pharm.ai

## Abstract

Accelerated development of lead structures is of high interest to the pharmaceutical industry in order to decrease development times and costs. We showcase how an intelligent combination of AI-based drug screening with state-of-the-art biophysics drives the rapid identification of novel inhibitor structures with high chemical diversity for cGMP-dependent 3’,5’-cyclic phosphodiesterase (PDE2). The starting point was an off-the-shelve chemical library of two million drug-like compounds. In a single *in silico* reduction step, we short-listed 125 compounds – the focused library – as potential binders to PDE2 and tested their binding behavior *in vitro* using MicroScale Thermophoresis (MST). Of this focused library, seven compounds indicated binding to PDE2, translating to a hit rate of 6%. Three of these compounds have affinities in the lower micromolar range. The compound with the highest affinity showed a *K*_*D*_ of 10 *µM* and is thus an excellent starting point for further medicinal chemistry optimization. The results show how innovative and structure-driven *in silico* approaches and biophysics can be used to accelerate drug discovery and to obtain new molecular scaffolds at a fraction of the costs and time – compared with standard high-throughput screening.

## Introduction

Phosphodiesterases are abundant enzymes, regulating cellular levels of the second messenger molecules cAMP and cGMP [1] and thus modulating regulatory pathways. Currently, twelve different families of phosphodiestrases are known with different 3D structures, kinetic properties, and modes of regulation [1]. Due to their broad biological roles, phosphodiesterases are attractive drug targets for different diseases states. Although the specific inhibition of phosphodiesterases via small molecules poses a challenge, sildenafil (Viagra) is one of the most popular, selective inhibitors for the phosphodiesterase family 5. Another selective inhibitor is the compound BAY60-7550 [2, 3].

cGMP-dependent 3’,5’-cyclic phosphodiesterase (PDE2), belonging to the phosphodiesterase family 2, is an enzyme with dual-specificity for cAMP and cGMP. It is encoded by the gene *PDE2A* in humans (Uniprot AC O00408). PDE2 hydrolyzes cAMP and cGMP, with its dual-specificity being determined by a freely rotating glutamine residue [4]. PDE2 plays an important role in growth and invasion of melanoma cells [5] and the development of specific inhibitors of PDE2 is an ongoing field of research [6, 7]. The selective inhibition of PDE2 was linked, for example, to improved object memory and synaptic plasticity [8].

Crystallographic structures of PDE2 are publicly available in the Protein Data Bank (PDB), rendering it a perfect target for structure-based drug discovery. Figure 1 depicts the crystal structure of PDE2 in complex with a selective inhibitor.

**Figure 1.**
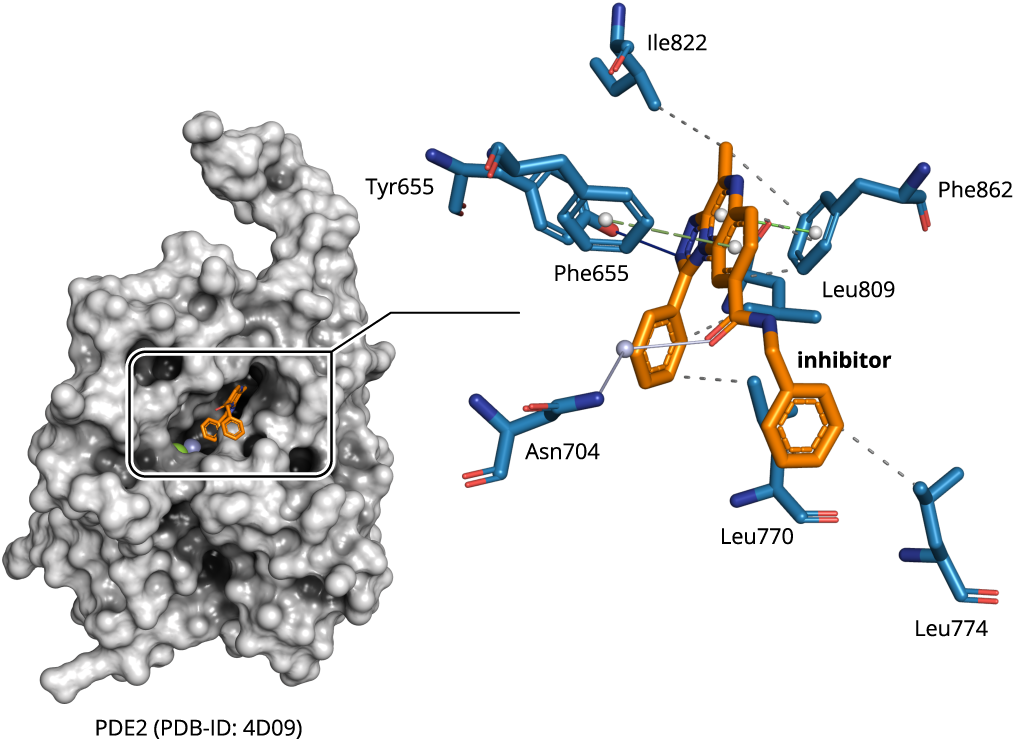
Crystallographic structure of PDE2 in complex with a selective inhibitor (PDB-ID: 4D09) [6]. The inhibitor forms characteristic, non-covalent protein-ligand interactions [9] that can be exploited to discover new small molecules with diverse scaffolds. Solid blue lines are hydrogen bonds, solid light blue lines are water-mediated hydrogen bonds, dashed green lines are *π*-stacking interactions, and dashed grey lines are hydrophobic contacts.

### Computational Drug Discovery

There is a plethora of approaches for computational drug discovery. However, the rapid *in silico* identification of small molecule ligands with high efficiency is still largely unaddressed. To answer the question, which approach might be the most effective in virtual screening (hit finding and target prediction), we were analyzing the reasons for drug promiscuity; a drug’s ability to bind to more than one target. It turned out that neither physicochemical properties of the ligands, nor their conformative flexibility fully explain why one small molecule is binding many targets while another one is selective [10]. However, we saw a clear signal when it came to similarities in the binding sites of the proteins to which a promiscuous drug binds. Similarities between the binding sites of otherwise unrelated (i.e. non-homologous) proteins allow a compound to bind to these different proteins [10]. This also holds true for similarities in the non-covalent interaction patterns of drug-target complexes [11].

Modularity is a design principle of biological systems, being inherent to proteins as well. Proteins achieve their functional diversity by shuffling domains [12, 13, 14], which again comprise small functional loops and peptides [15, 16]. This implies a limited diversity in protein structures, which is expressed in the saturated fold space. Similar observations for ligand binding sites in proteins [17, 18, 19] suggest that diversity in binding sites could be limited as well. The established assumption of binding site diversity – originating in the concept of single-target drugs – has to be reconsidered. Instead, the high modularity of proteins on each level – also holding true for binding sites and protein-protein interfaces [20], suggesting a degenerated pocket space to be rooted in protein evolution [21, 22].

Recently, there is a growing trend towards ultra-high throughput pipelines for computational drug discovery. Examples include the identification of novel *β*-lactamase inhibitors and dopamine receptor agonists [23], melatonin receptor ligands [24], or protein-protein interaction inhibitors [25]. Even though these results are exiting and promising, they come with a major restriction. All of the aforementioned methods depend on computation-intensive docking protocols and molecular dynamics simulations, or combinations thereof. A “brute force”-like approach is applied to collections of millions, or even billions, of compounds, which is infeasible for the average early stage drug discovery project with limited resources and timelines. Generative chemistry, driven by artificial intelligence [26], might be more efficient but often lacks the desired novelty of the generated small molecules as these methods depend on training data of known chemical entities.

### Focused Library Service

To this end, we developed a software suite, the PharmAI *DiscoveryEngine*, turning our findings about the modularity of ligand binding sites into a fully integrated virtual screening engine that allows for effective and efficient hit finding for a given drug target. In contrast to the aforementioned pipelines, our technology differentiates as follows:

▸ The knowledge-based *DiscoveryEngine* exploits our findings about modularity of ligand binding.
▸ It features the prediction of diverse chemical scaf-folds.
▸ The mapping of chemical scaffolds to an arbitrary chemical library to pinpoint the important subsets with high hit probability, i.e. the *Focused Library*.

By following this intelligent selection strategy, computational intensive tasks, such as docking or molecular dynamics simulation, might only be conducted as a subsequent refinement step. This concept allows for an improvement in prediction quality at a tiny fraction of costs and resources. Figure 2 illustrates the workflow and timeline of a typical *Focused Library* screening project – bringing together PharmAI’s Focused Library Service and 2bind’s MicroScale Thermophoresis Service to leverage each technology’s potential.

**Figure 2.**
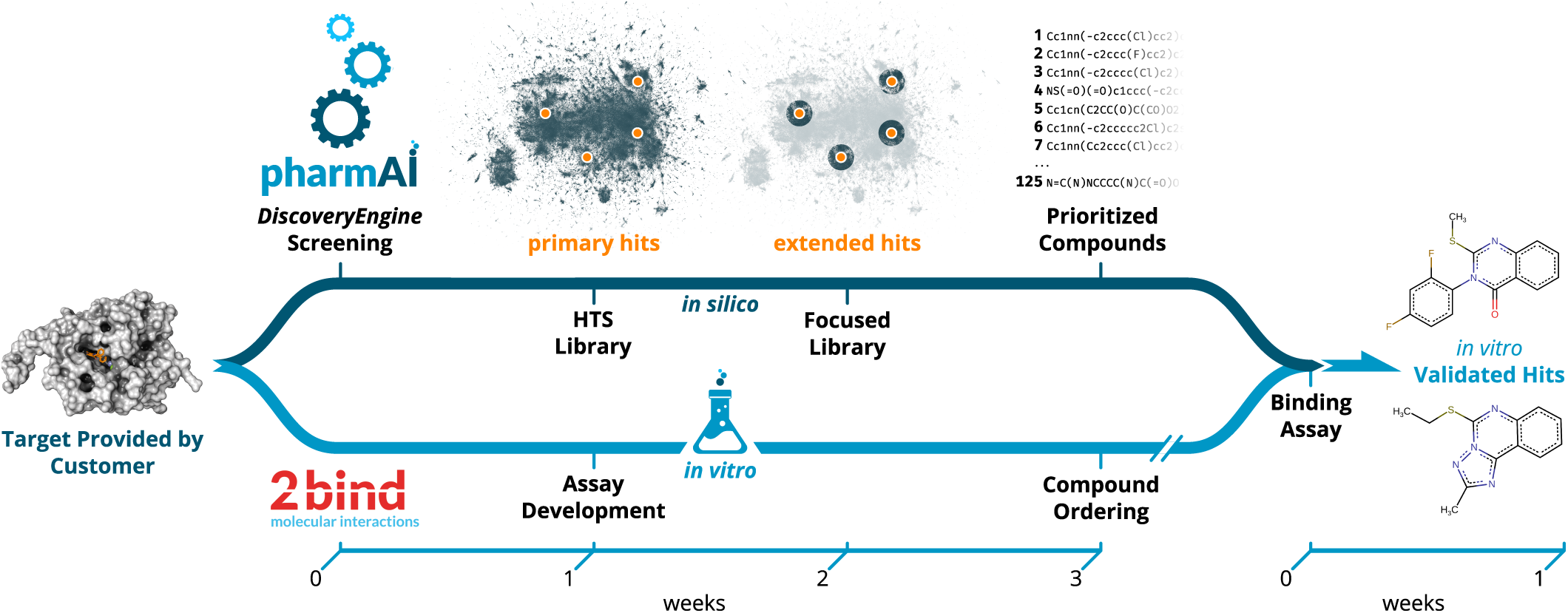
Workflow of the PharmAI·2bind *Focused Library Service*. Upon the definition of the target by the customer, the *in silico* and *in vitro* pipelines are launched in parallel. While the PharmAI *DiscoveryEngine* predicts primary hits with diverse scaffolds tailored to the given target, the assay development for downstream validation of the predictions starts at 2bind. The predicted primary hits (orange dots) are mapped to the desired HTS library (dark blue background). Based on the primary hits, compounds are selected from the HTS library via a sophisticated and optimized chemical similarity screening. Upon completion of the compound prioritization, the compounds are ordered from the vendor. With the proper binding assay in place, compounds can be immediately tested *in vitro*. As a final result, the customer receives *in vitro* validated hits. The whole process can be completed within four weeks, excluding protein sourcing and compound procurement.

### MicroScale Thermophoresis (MST)

Advances in computational pre-selection of large compound libraries have to be met with adequate, state-of-the-art *in vitro* methods for hit validation. Moreover, consumption of often precious protein and compound material has to be taken into account. A method that combines speed and efficiency with ultra-low sample consumption and that can deliver the important steady-state affinity (*K*_*D*_) in high quality for each pre-selected library compound is MicroScale Thermophoresis (MST). From a technical perspective, this method is fully compatible with classical 384-well plate compound library formats and can be combined with lab automation (e.g. LabCyte Echo acoustics, contact-less compound preparation) to achieve quick results. The high sensitivity and wide applicability of MST (proteins, DNA, RNA, lipids, compounds, fragments, particles) relies on its unique measurement principle (Figure 3a). For MST, the target molecule is labeled with a special fluorescent dye. A general property of fluorescence is that its intensity decreases with increasing temperature. This effect is called temperature-related intensity change (TRIC). Now this fluorescence change can be further manipulated by the binding of a ligand to the labeled target molecule (binding in close proximity to the dye or via conformational changes of the target). Together with an additional readout principle, thermophoresis, the MST signal is sensitively affected by various different molecular parameters, which usually all change upon binding of a ligand to a target molecule. The output of MST are classical and well-described ligand-binding dose-response curves, which can be fitted to yield the steady-state affinity (*K*_*D*_) of an interaction (Figure 3b).

**Figure 3.**
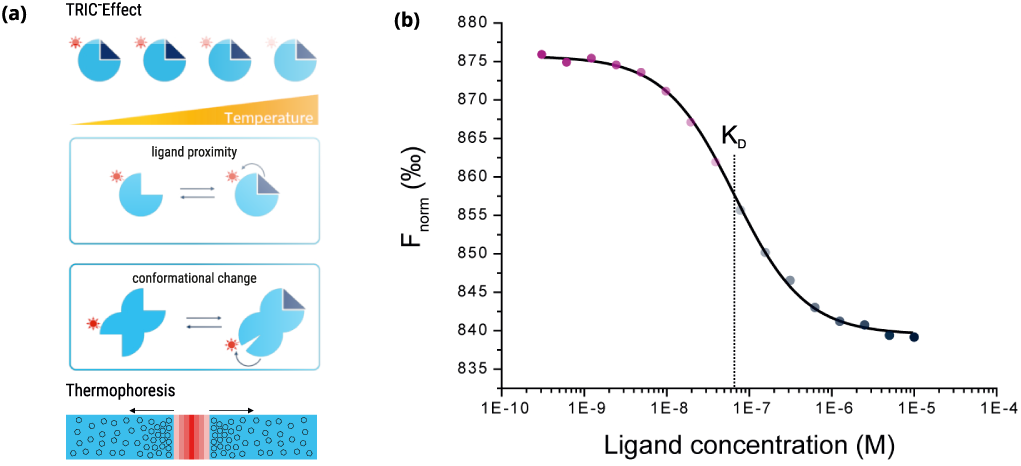
Principle and output of the MST technology. (a) Schematic illustration of the TRIC effect (top and middle) and thermophoretic movement of molecules in a temperature gradient (bottom). (b) The calculated *F*_norm_ values from the MST traces are dose-dependent and can be well described by the law of mass action. A fit of these values returns the dissociation constant *K*_*D*_ of the interaction.

## Results and Discussion

In this case study, we applied the full *Focused Library Service* pipeline to predict new lead structures from a highly diverse compound collection. We prioritized 125 compounds from a screening collection that contains approximately 1.9 million compounds, corresponding to only ≈ 0.0066% of the original library size. Out of the 125 compounds tested, seven molecules indicated binding to the target in an MST assay, which translates to an exceptional high hit rate of 5.60%.

For each of the identified compounds, we performed a chemical similarity search in the ChEMBL [27] and BindingDB [28] databases. These databases contain a comprehensive set of experimentally validated activities of small molecules against drug targets and allow to assess the chemical novelty of the predicted hits in comparison to known molecules binding to the same target. The identified binders along with the closest counterparts in ChEMBL [27] and BindingDB [28] are listed in Table 1 and sorted according to the ranks of primary hits predicted by the PharmAI *DiscoveryEngine*.

**Table 1.**
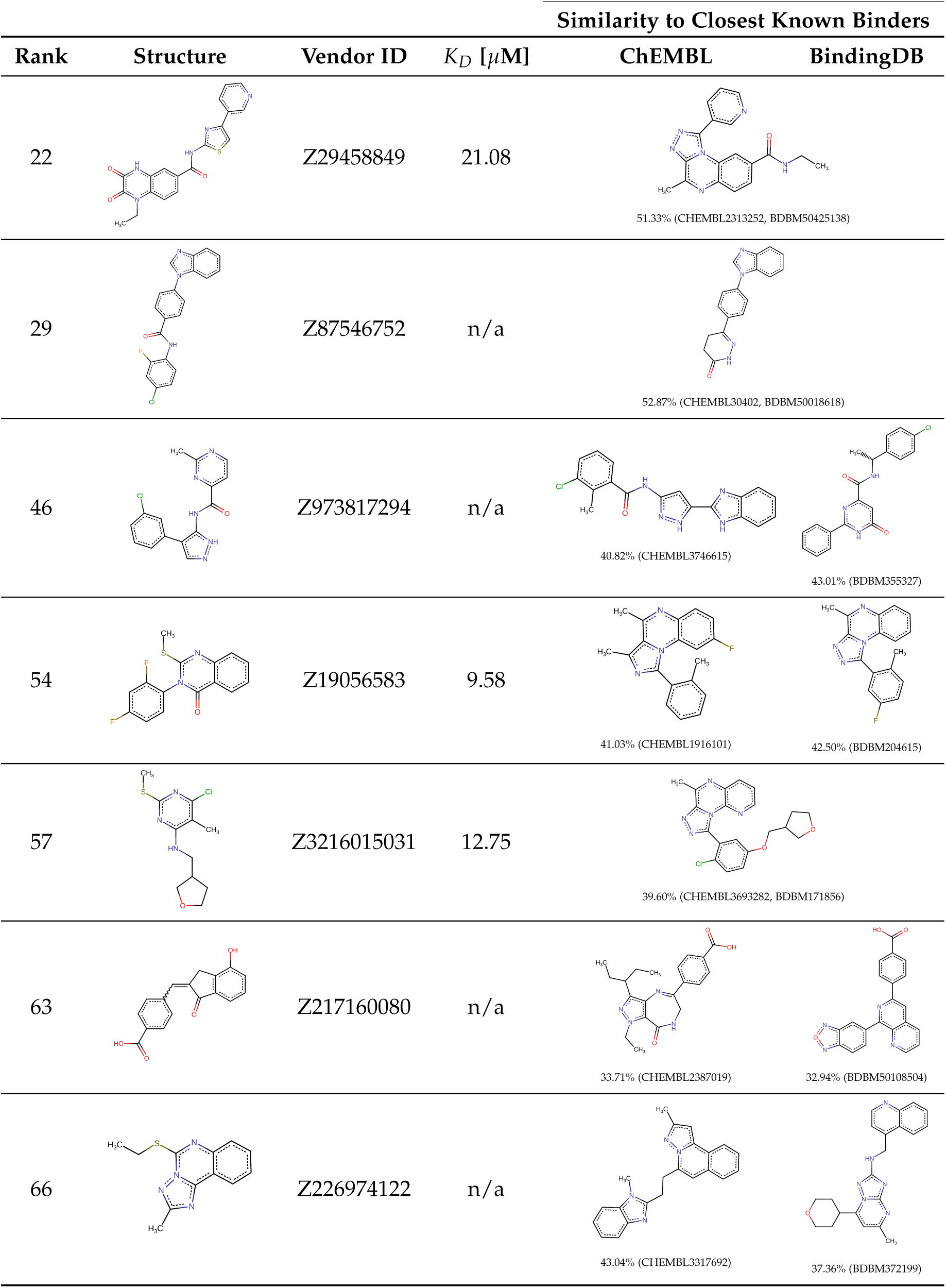
Identified Enamine compounds with indicated binding to PDE2A. For each compound the closest entry in ChEMBL [27] and BindingDB [28] that is associated to PDE2 is shown. Compounds are ordered according to the rank of primary hits predicted by the PharmAI *DiscoveryEngine*.

| Rank | Structure | Vendor ID | $K_D$ [μM] | Similarity to Closest Known Binders | |
| --- | --- | --- | --- | --- | --- |
|  |  |  |  | ChEMBL | BindingDB |
| 22 |  | Z29458849 | 21.08 | 51.33% (CHEMBL2313252, BDBM50425138) |  |
| 29 |  | Z87546752 | n/a | 52.87% (CHEMBL30402, BDBM50018618) |  |
| 46 |  | Z973817294 | n/a | 40.82% (CHEMBL3746615) | 43.01% (BDBM355327) |
| 54 |  | Z19056583 | 9.58 | 41.03% (CHEMBL1916101) | 42.50% (BDBM204615) |
| 57 |  | Z3216015031 | 12.75 | 39.60% (CHEMBL3693282, BDBM171856) |  |
| 63 |  | Z217160080 | n/a | 33.71% (CHEMBL2387019) | 32.94% (BDBM50108504) |
| 66 |  | Z226974122 | n/a | 43.04% (CHEMBL3317692) | 37.36% (BDBM372199) |

### Validation of Predicted Compounds

All 125 predicted compounds were validated using a 2bind MST screening assay and docked to the PDE2 target using AutoDock Smina with a flexible docking protocol [29, 30]. For three of the seven identified binders – Z19056583, Z3216015031, and Z29458849 – *K*_*D*_ values could be determined (10.15, 14.24, and 24.91 *µ*M, respectively); response amplitudes and fit signal/noise ratios met the quality requirements in all cases. Figure 4a shows dose-response data and calculated docking poses. The other four hits were weak binders with apparent *K*_*D*_ values above the maximum screening concentration of 100 *µ*M (Figure 4b). No ligand-related artifacts, such as compound-induced sticking, aggregation, or fluorescence effects, were observed. The reference protein used for docking calculations is shown in Figure 4C. Docking scores with the default AutoDock Smina scoring function were in the range between -12.0 and -7.4 kcal·mol^−1^. The control compound BAY 60-7550 achieved a score of -10.7 kcal·mol^−1^ when docked to the reference structure. For each of the predicted compounds, the top-scoring docking poses were analyzed regarding their non-covalent interactions [9] with the target binding site. Phe862 is involved in *π*-stacking interactions for each of the poses and seems to be able to form sandwich *π*-stacking with the help of Phe655. In general, the interactions are driven by hydrophobic contacts, stacking interactions, and halogen bonds.

**Figure 4.**
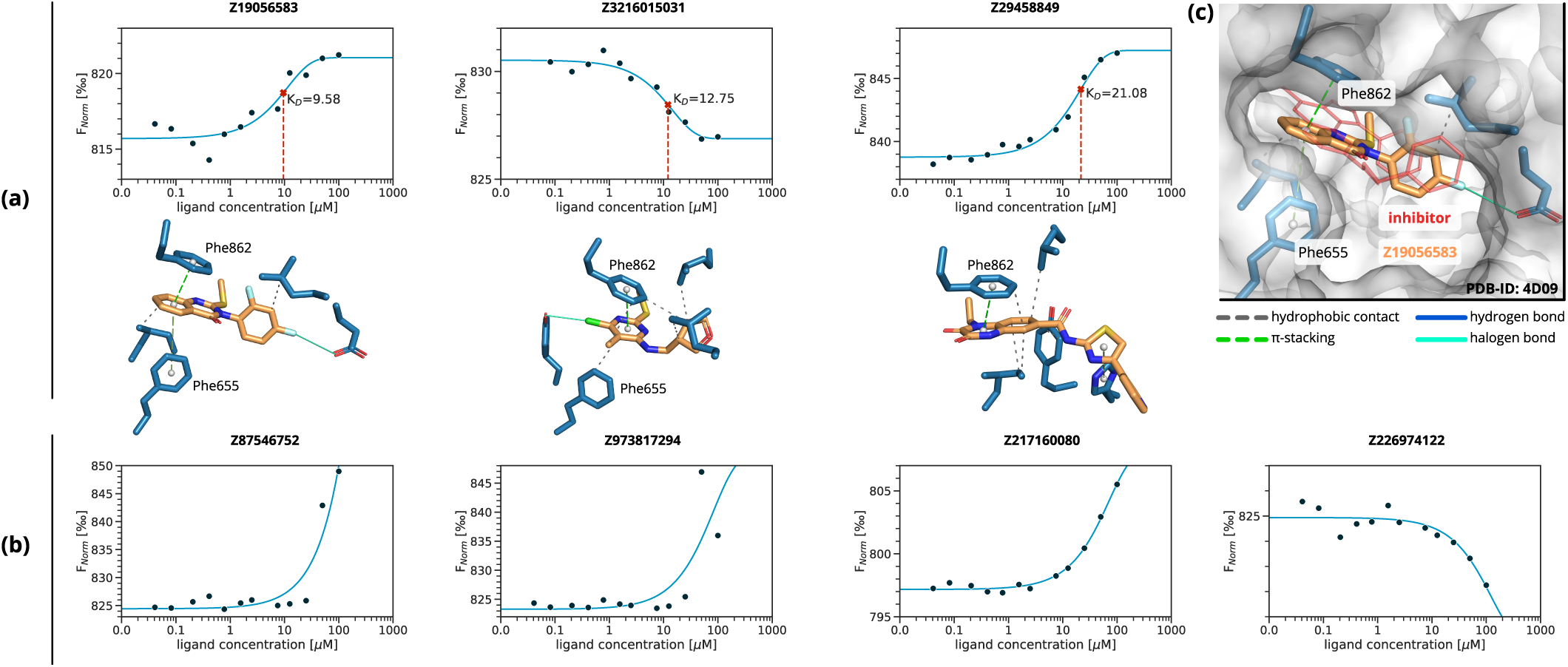
MST dose-response graphs and docking poses for the seven identified binders. Non-covalent interactions between the predicted compounds and PDE2 were calculated with PLIP [9]. Solid cyan lines are halogen bonds, dashed green lines are *π*-stacking interactions, and dashed grey lines are hydrophobic contacts. (a) Compounds for which a full dose-response curve could be obtained. The solid line is a *K*_*D*_ 1:1 fit, the *K*_*D*_ value is indicated by the dashed vertical line. (b) Compounds with weak binding and apparent *K*_*D*_ values greater than the maximum screening concentration of 100 *µ*M. Note that the solid line is no *K*_*D*_ fit, but rather serves as a visual guide for the observed increase in Fnorm values at the higher ligand concentrations. (c) Docking pose of Z19056583 to the reference structure of PDE2 (PDB-ID: 4D09) [6]. The reference ligand, used to define the binding site, is shown in red line representation.

### Hit Rate

We observe an exceptionally high hit rate of 5.60%. If only compounds with measured *K*_*D*_ values are taken into account, the hit rate is 2.40%. A benchmark study by Chiba *et al*. [31], where virtual screening predictions of 16 scientific groups were combined and assessed *in vitro*, achieved a hit rate of 0.22% for the NAD-dependent deacetylase sirtuin-1. Based on these findings, with the results presented in this case study, the PharmAI *DiscoveryEngine* shows an 11- to 25-fold improvement over existing technologies. Simultaneously, the number of compounds needed to be screened was 25 times lower (125 vs. 3192), while still achieving the same number of seven hits [31]. In a traditional HTS setup with a hit rate of around 0.02% [32], *in vitro* testing of 35,000 compounds would have been necessary to achieve a comparable number of hits. This would translate to a tremendous increase in costs and time and highlights the strengths of efficient combination of *in silico* driven compound selection and fast testing via MST.

### Diversity of Predictions

A two-dimensional representation, generated with the Uniform Manifold Approximation and Projection for Dimension Reduction (UMAP) algorithm [33], allows to relate the predictions of the PharmAI *DiscoveryEngine* to the full HTS library. Each compound is hereby represented by an optimized, high-dimensional chemical fingerprint. It is evident that the predictions of the *DiscoveryEngine* hit the full diversity of the library (Figure 5a) and are distributed across the chemical space. At the same time, the compounds that indicate binding to PDE2 are of high scaffold diversity (Figure 5b) and located it different areas of the chemical space.

**Figure 5.**
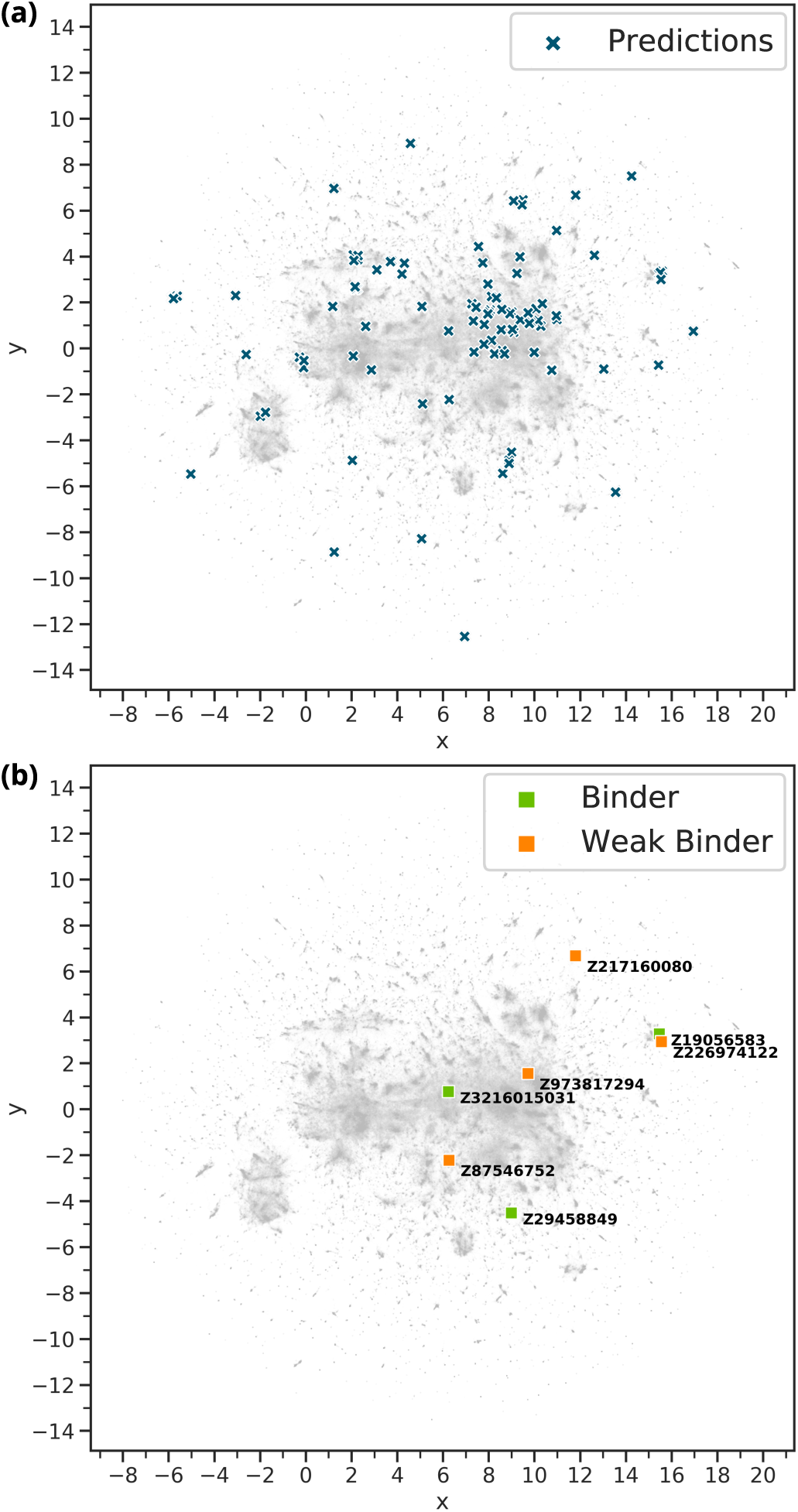
Relation of the *DiscoveryEngine* predictions to the Enamine HTS library. Each grey pixel represents one of the 1.9 million compounds of the library. The two-dimensional representation was generated with UMAP [33]. (a) The 125 predicted compounds (blue crosses) in relation to the HTS library. They are distributed equally over the whole chemical space. (b) The compounds with indicated binding to PDE2 in relation to the HTS library.

### Similarity to Known Compounds

As well as traditional HTS techniques, computational drug discovery faces the hurdle to discover new chemical entities. Ideally, the identified compounds should be new chemical species with scaffolds divergent to known compounds. This allows to circumvent patent protection and to target new molecular mechanisms. While generative approaches [26] often have difficulties addressing this issue, the predictions made by the PharmAI *DiscoveryEngine* show a high scaffold diversity and low similarity to known PDE2 inhibitors. The closest compound to our predictions shows a similarity of 52.87% (CHEMBL30402) and was measured with an IC50 activity against PDE2 of 34 *µ*M (CHEMBL759152). On average, the similarity to compounds with known activity against PDE2 is 43.20% for ChEMBL and 42.80% for BindingDB. These numbers emphasize that the scaffold diversity of our predictions is high and new lead structures for PDE2 inhibition were obtained. Selectivity can be an issue for phosphodiesterase inhibitors, e.g. caffeine is known to be an unselective inhibitor [1]. By providing diverse chemical scaffolds as starting points for further development, the downstream risk of undesired off-target hits is reduced.

## Conclusions

We showcased an application of the PharmAI *Focused Library Service* in combination with the 2bind biophysical hit-validation. The application of our technology enabled the quick identification of seven small molecules in a library of 1.9 million compounds that indicate binding to PDE2. Due to the intelligent and knowledge-based prediction of diverse molecular scaffolds, the selection of only 125 compounds from the HTS library, and rapid and economical validation by MST, screening costs and timelines are drastically reduced. Our method achieves a hit rate of 2.40% if compounds with measured *K*_*D*_ values are taken into account, and 5.60% if weak binders are considered. This translates to an 11- and 25-fold improvement, respectively, over other virtual screening techniques [31].

## Methods

### Generation of the Focused Library

The PharmAI *DiscoveryEngine* is a knowledge-based *in silico* drug screening platform optimized for the processing of 3D protein structures. The entry point to a *DiscoveryEngine* screening is the drug target – normally one or more crystal- or NMR structure of the protein – in complex with its ligand(s). These complexes may come directly from the PDB, from proprietary repositories, or from simulations such as docking, molecular dynamics, and homology modeling.

Initially, the *DiscoveryEngine* analyzed the PDE2 binding site for geometrical properties, non-covalent interaction patterns, and physico-chemical properties in 36 PDB structures (including PDB structures 4HTX and 5U7D binding BAY-60-755). In a next step, these properties were converted into profiles representing the target’s binding site and the way it interacts with ligands. Subsequently, the derived profiles were used to screen PharmAI’s data warehouse of preprocessed protein and small molecule data. As a result, a primary hit list of 54 scaffolds with a good fit for the target binding site was returned.

These scaffolds are then mapped to the chemical library by advanced chemical similarity. For each of the mapped scaffolds, nearest neighbors in the chemical library were queried and ranked by significance. Enamine’s HTS Collection^a^ as of June 19, 2019 was used as chemical library. At this time, the collection contained 1,963,425 compounds. For all compounds, chemical similarities were computed using Extended-Connectivity Fingerprints [34] with highly optimized parameters. Dimension reduction for visualization of the chemical library was conducted with UMAP [33]. ChEMBL version 26 and BindingDB as of 19 March 2020 were used to identify existing inhibitors. Entries in these databases mapped to UniProt AC O00408 were considered to be potentially active against PDE2, irrespective of inhibitory concentrations.

### *In Vitro* Validation

The PDE2 protein was purchased from Antibodies Online (13031-H08B/ABIN2005384, lot LC10NO1404) and labeled with a 647 *nm*, red-fluorescent MST dye (NHS coupling chemistry). The predicted compounds were purchased from Enamine as 10 *mM* DMSO stocks. 12-concentration compound serial dilutions were prepared via contactless, high-precision, acoustic dilution (LabCyte Echo platform). The MST screening was performed on an NT. Automated instrument with a “pico-red” detector setting and “medium” MST-power setting. The compound screen was validated using BAY 60-7550, a potent and selective PDE2 inhibitor with a described *K*_*i*_ of 3.8 *nM* [3], which was purchased from Sigma-Aldrich (SML2311, lot 0000047650) as a powder stock and solved to 10 *mM* in water-free DMSO. The control was analyzed before and after the 125-compound set and the aggregated determined *K*_*D*_ value was 33.0 *nM*. MST screening data were exported with MO.ScreeningAnalysis (1.0.3, NanoTemper Technologies) using an MST on-time interval of 4-5 seconds and otherwise standard settings and further analyzed with 2bind screening-software tools.

## Footnotes

a enamine.net/hit-finding/compound-collections/screening-collection/hts-collection

